# Expanding genetic code to generate human brain organoids with both vasculature and microglia

**DOI:** 10.64898/2026.07.08.737383

**Authors:** Haishuang Lin, Yaqing Wang, Hu Du, Yuxin Qin, Haoyu Zhang, Peng Wang, Lin Wei, Jianhua Qin

## Abstract

Brain organoids offer an invaluable model system for studying human brain development and disease. However, the establishment of high-fidelity brain organoids with multiple cell lineages including vasculature and immune cells remains a huge challenge. Here, we present a new strategy to generate human cerebral organoids with vasculature and microglia-like cells using genetic code expansion technology (GCE-T) via site-specific protein engineering. The strategy integrates orthogonal genetic translation machinery in hPSCs via PiggyBac transposon system, enabling temporally control of ETV2 expression and endothelial differentiation in hPSC-derived cerebral organoids. The vascularized human cerebral organoids (vhCOs) exhibit coordinated development of multiple cell lineages and blood-brain barrier (BBB) features. Moreover, vhCOs form perfusable vascular network after transplanted in the immune-deficient mice. Single-nucleus RNA sequencing reveals enhanced neurovascular interactions, multi-brain-regional identities, diverse neuronal subtypes and specialized endothelial subclusters in vhCOs, closely resembling human fetal brain. Strikingly, we identify enriched microglia-like cells comprising three distinct subtypes in vhCOs, which contribute to microglia-vascular interactions and synergistically modulate vascular development. Upon Zika virus (ZIKV) infection, vhCOs show neurovascular dysfunction and impaired microglia development, offering new insights into viral-induced neurodevelopmental disorders. This study offers a unique platform for producing more valuable brain organoids with vasculature and immune components, opening a new avenue to advance organoid research and applications.

## Introduction

Brain organoid derived from human pluripotent stem cells (hPSCs) is the most extensively studied human organoids in recent years. It can recapitulate key features of the early or even mid-prenatal brain, which has been utilized for studying brain development and neurologic disorders^[1–3]^. *In vivo,* brain organogenesis is characterized by a complex process that are temporally regulated by the multicellular types, including neurons, vascular cells, astrocytes and resident immune cells^[4,5]^. Specially, vascular network is critical for supporting neurodevelopment, multiple brain region formation and functional maturity^[6,7]^. Immune cells, such as microglia, play a crucial role in immune surveillance while regulating brain development, neuronal survival, synapse formation and elimination of apoptotic cells^[8,9]^. Multicellular interactions are pivotal to maintaining homeostasis within the brain microenvironment. Currently, several approaches have been developed to generate brain organoids integrated with vascular cells, such as co-culture^[10,11]^, organoid fusion^[12]^, and genetic induction^[13]^. However, establishing high-fidelity brain organoids with multiple cell lineages including vasculature and immune components remains a significant challenge.

Genetic code expansion technology (GCE-T) is a powerful platform to precisely engineer protein structure and function. It employs a reprogrammed tRNA/aminoacyl-tRNA synthetase (aaRS) pair and a repurposed nonsense codon (e.g., TAG, TGA, or TAA) to achieve site-specific incorporation of noncanonical amino acids (ncAAs)^[14]^. The incorporation of ncAAs can be functionalized with fluorescent probes, metal-chelating groups, photocrosslinkers, or post-translational modifications in prokaryotic and eukaryotic cells^[15]^. The orthogonal aaRS/tRNA pair specifically recognizes the desired ncAA in response to the nonsense codon, without cross-reactivity with endogenous translation systems. Especially, the Methanosarcina mazei (MmPyl) aaRS/tRNA pairs have been widely utilized in mammalian cells^[16]^. However, this GCE-T has not been explored in 3D tissues or organoids.

In this work, we made the first attempt to generate human cerebral organoids with functional vasculature and microglia-like cells derived from hPSCs using GCE-T, which is designed to be transcriptionally “on” but translationally “off” without the inducer. Our system enables precise translational control, exhibiting minimal leakage and rapid responsiveness. We integrated the orthogonal genetic translation machinery (GTM) into hPSCs via site-specific incorporation of a ncAA at the TAG site, enabling temporally control of ETV2 expression and endothelial differentiation in hPSC-derived cerebral organoids. Our data demonstrated that the engineered organoids recapitulate key aspects of human fetal brain development by integrating functional vasculature and blood-brain barrier (BBB) features. We found that these cerebral organoids possess enriched microglia-like cells that could contribute to vascular development. Single-nucleus RNA sequencing reveals a strong correlation between the transcriptional signatures of cerebral organoids and those of the human fetal brain. We further leveraged this novel organoid platform to model Zika virus infection, elucidating previously unreported pathophysiology such as neurovascular dysfunction and impaired microglial development.

## Results

### Construction of bio-orthogonal genetic translation machinery in hPSCs

In order to generate brain organoids with multicellular lineages by site-specific protein engineering, we initially attempted to control endothelial differentiation in hPSCs using bio-orthogonal GTM. ETV2, a key transcriptional regulator of endothelial cell (EC) fate, can directly reprogram somatic cells (e.g., fibroblasts) to ECs when transiently expressed^[17–19]^. To achieve full-length ETV2 expression in human pluripotent stem cells (hPSCs), we established stable H9^GTM^ and iPS^GTM^ cell lines by integrating GTM into hPSCs via the PiggyBac transposon system. This GTM comprises the Methanosarcina mazei pyrrolysyl-tRNA synthetase/tRNA (MmPyIRS/tRNA) pair and an *ETV2* gene containing a premature termination codon (PTC) at residue A122. The orthogonal translation system enables site-specific incorporation of a ncAA Nε-2-azidoethyloxycarbonyl-L-lysine (NAEK) at the PTC site, yielding full-length ETV2 protein (**Fig. 1a, b**). Notably, H9^GTM^ showed marked *ETV2* expression only in the presence of NAEK (**Fig. 1c**), indicating the efficient suppression of stop codon by the MmPyIRS/tRNA system. Engineered H9^GTM^ cells exhibited compact colony morphology from passage 1 to passage 20 (**Fig. 1d, and Supplementary Fig. 1a**). Additionally, quantitative real-time PCR (qRT-PCR) analysis revealed that H9^GTM^ cells showed sustained expression of pluripotent markers (*OCT4, NANOG*) and elevated expression of *ETV2, MmPylRS, and tRNA/18S* compared to H9 controls (**Fig. 1e**). Immunostaining also confirmed expression of pluripotency markers (OCT3/4, NANOG, SSEA4) and ALP (**Supplementary Fig. 1b**). Engineered H9^GTM^ cells exhibited proper proliferation and survival (**Supplementary Fig. 1c, d**). Sanger sequencing confirmed correct PTC integration and genomic stability over 20 passages without reverse mutation (**Fig. 1f**). H9^GTM^ can differentiate into cells of all three germ layers in embryoid bodies (EBs), including NESTIN^+^ ectoderm, Branchyury+ mesoderm and FOXA2^+^ endoderm (**Supplementary Fig. 1e**). They further formed teratomas containing structures representative of each germ layer (e.g. neural rosettes, muscle and gut-like epithelium), confirming the pluripotency of H9^GTM^ cells (**Fig. 1g**). These results demonstrate successful integration of bio-orthogonal GTM in hPSCs.

**Figure 1.**
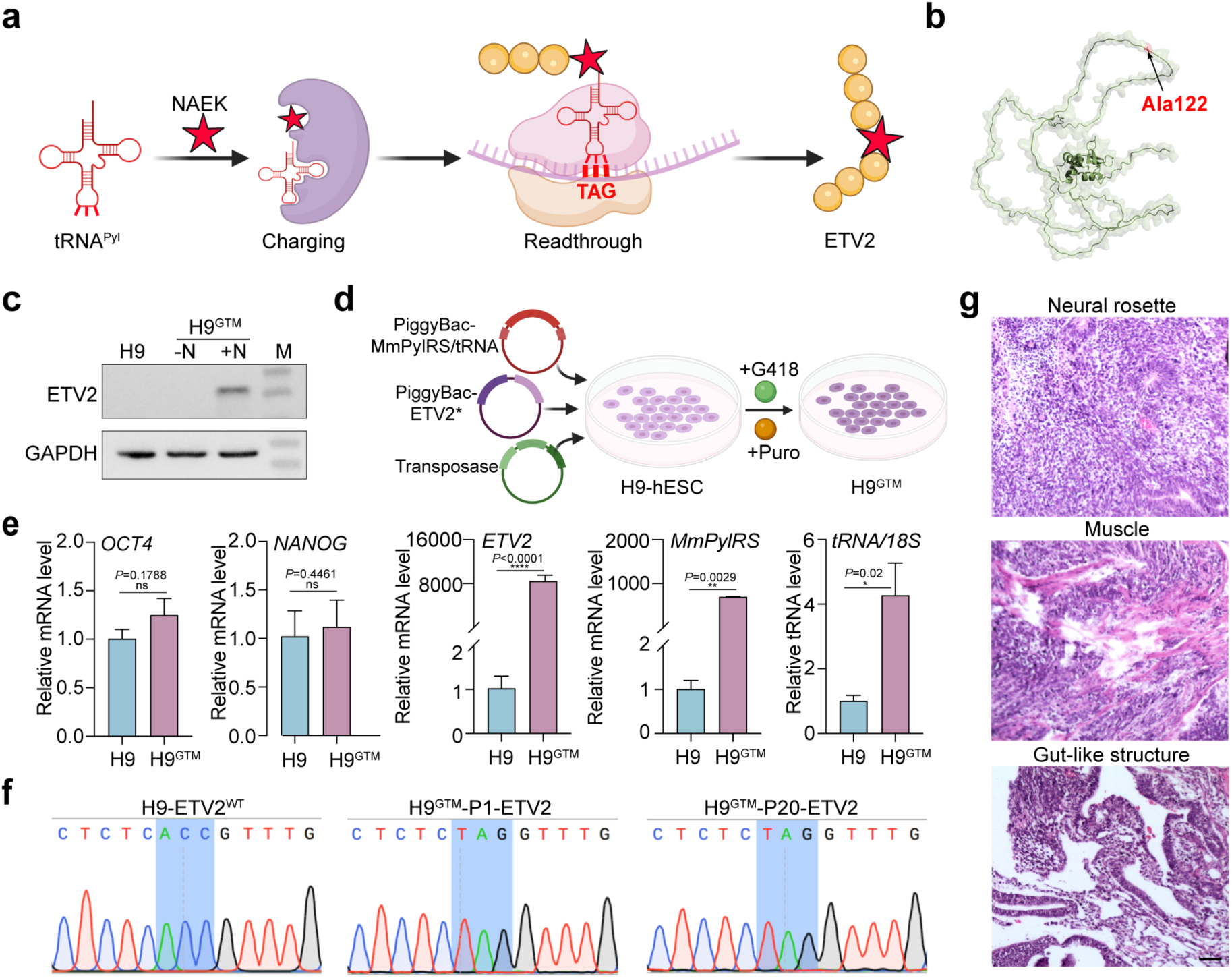
Establishment of bio-orthogonal genetic translation machinery (GTMs) in hPSCs. (**a**) Schematic of the genetic translation system. A premature termination codon (PTC) was introduced into the human *ETV2* gene. The *Methanosarcina mazei pyrrolysyl* (MmPyl)/tRNA_CUA_ translational machinery enables site-specific incorporation of a non-canonical amino acid (ncAA) at the PTC, allowing temporally controlled expression of full-length ETV2 protein. (**b**) Structural illustration showing the PTC introduction site (A122 residue, highlighted in red). The structure was predicted using AlphaFold2. (**c**) Western blot analysis of ETV2 expression following NAEK treatment in H9 cells harboring the GTM system. Data are representative of three independent experiments (n = 3). (**d**) Schematic of the screening strategy for stable H9^GTM^ lines. Asterisk indicates the introduced PTC in ETV2. (**e**) Quantitative real-time PCR analysis of pluripotency markers (*OCT4, NANOG*), *ETV2, MmPylRS,* and *tRNA_CUA_*. Data are shown as mean ± SD (n = 3 from three independent batches). (**f**) Sanger sequencing confirming the correct mutation site and the absence of reverse mutation after 20 passages. (**g**) Teratoma assay demonstrating differentiation of H9^GTM^ into three germ layer tissues. Scale bar, 200 µm.

To further validate the ability of endothelial differentiation in engineered hPSCs with an expanded genetic code, we incubated H9^GTM^ cells with NAEK for two days in 2D cultures (**Supplementary Fig. 2a**). Crucially, a portion of EC-like cells were observed in H9^GTM^ cells after administration of NAEK (**Supplementary Fig. 2b**), accompanied by upregulated gene expressions of *ETV2* and *VE-Cadherin* (**Supplementary Fig. 2c**). These data indicated that full-length ETV2 was produced depending on NAEK at the translational level and the functional ETV2 protein could regulate the rapid differentiation of ECs from H9^GTM^ cells.

### Generation of human cerebral organoids with vasculature, microglia and BBB features via GCE-T

In principle, integration of orthogonal GTM in hPSCs allows temporally control of ETV2 expression in their derived cells and complex tissues. Since hPSCs can self-organize into brain organoids in 3D culture^[20]^, we sought to generate cerebral organoids containing vasculature from H9^GTM^ by inducing ETV2 expression via stop codon suppression during development. H9^GTM^ cell suspension was seeded in concave wells to form EBs, followed by embedded in Matrigel for neuroepithelial expansion (**Fig. 2a**). These spheroids were then transferred to spinning bioreactors for suspended culture. On day 18, we introduced NAEK to drive ETV2 expression and initiate vascularization of organoids, thereby generating vascularized human cerebral organoids (referred as vhCOs) (**Fig. 2a**). At this stage, organoids have formed early neural structures but not yet matured, offering a favorable time point for endothelial cell induction to support vascular-neural development. Furthermore, we found that the morphology of vhCOs was significantly affected by day 6 of NAEK treatment compared to hCO (**Supplementary Fig. 3**). Notably, H9^GTM^-derived vhCOs exhibited progressive growth and larger size than H9-derived hCOs, reaching ∼4.5 mm diameters by day 180 (**Fig. 2b, c**). Immunostaining of 30-day vhCOs showed the presence of SOX2^+^ neural progenitor cells, TUJ1^+^ neurons, CD31^+^ ECs (**Fig. 2d**), vascular lumen-like structure (**Fig. 2e**), and COL4^+^/CD31^+^ vascular basement membrane (**Fig. 2f, and Supplementary Fig. 4a**). These data demonstrated the endothelial differentiation and vasculature construction in organoids during early developmental stages.

**Figure 2.**
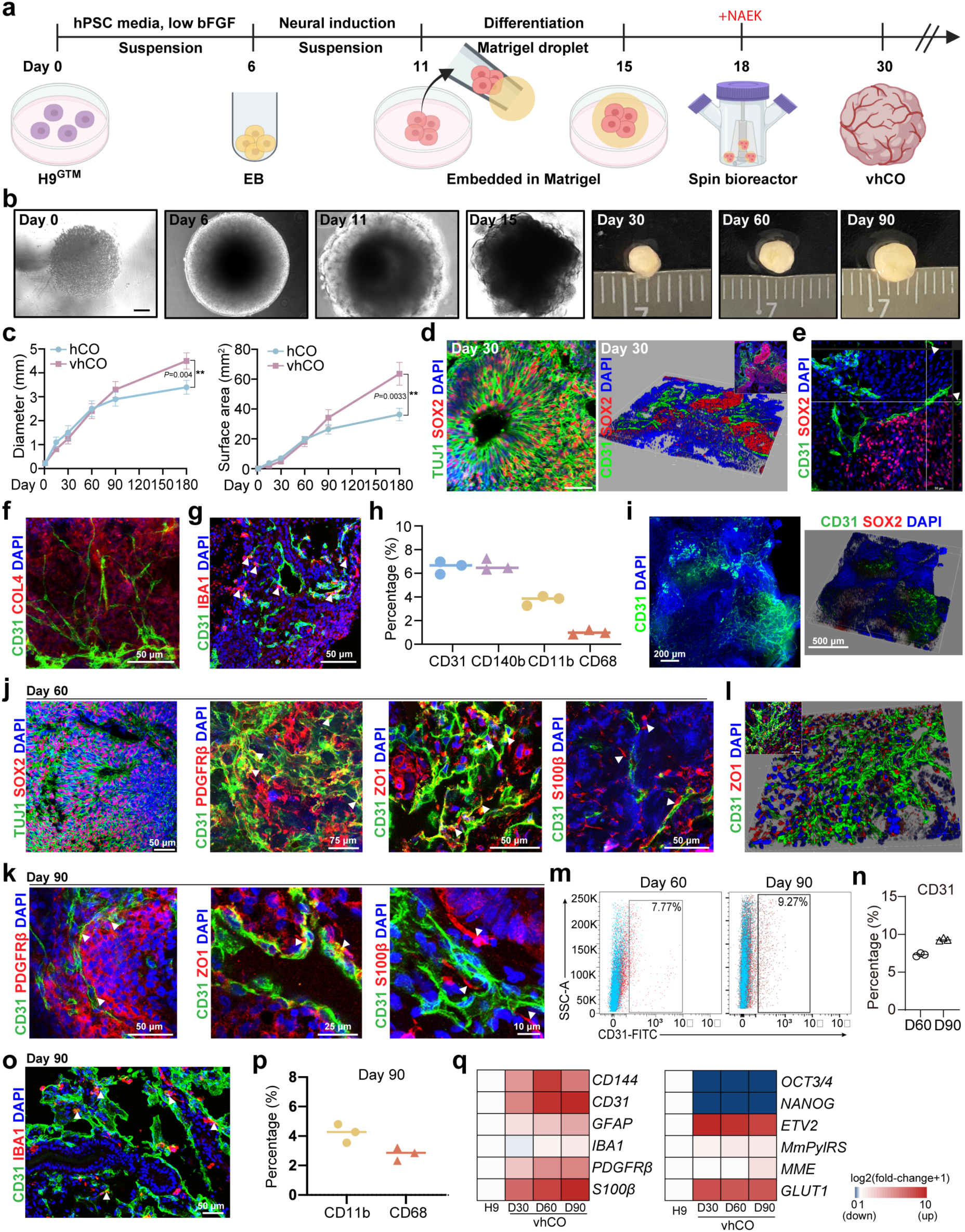
Generation of vascularized human cerebral organoids via genetic code expansion. (**a**) Schematic of brain organoid differentiation and ncAA treatment at day 18. (**b**) Bright-field images of organoids at indicated developmental stages. Scale bars, 100 µm. (**c**) Quantitative analysis of organoid size in hCOs and vhCOs over time. Data are shown as mean ± SD (n = 3 from three independent batches). (**d**) Immunostaining of SOX2+/TUJ1+ (left) and CD31+/SOX2+ (right) cells in day-30 organoids. Scale bars, 100 µm (left) and 50 µm (right). (**e**) Orthogonal section of a day-30 vhCO showing lumen-like structures (white arrows). (**f, g**) Immunostaining of COL4+/CD31+ (**f**) and CD31+/IBA1+ (**g**) cells in day-30 organoids. Scale bars, 100 µm (left) and 50 µm (right). (**h**) Flow cytometry analysis of CD31, CD140b, CD11b, and CD68 in day-30 organoids. Data are shown as mean ± SD (n = 3 from three independent batches). (**i**) Optical tissue clearing of a day-30 vhCO revealing the vascular network. Scale bars, 200 µm (left) and 500 µm (right). (**j, k**) Immunostaining of SOX2+/TUJ1+, CD31+/PDGFRβ+, CD31+/ZO1+, and CD31+/ S100β+ cells in day-60 (**j**) and day-90 (**k**) organoids. Scale bars, 100 µm (left) and 50 µm (right). (**l**) Confocal 3D reconstruction of CD31 and ZO-1 immunostaining showing a vascular lumen-like structure in a day-90 vhCO. Scale bar, 50 µm. (**m, n**) Flow cytometry analysis of CD31+ cells in day-60 (m) and day-90 (n) organoids. Data are shown as mean ± SD (n = 3 from three independent batches). (**o**) Immunostaining of CD31+/IBA1+ cells in day-90 vhCOs. Scale bars, 100 µm (left) and 50 µm (right). (**p**) Flow cytometry analysis of microglial markers CD11b and CD68 in day-90 organoids. Data are shown as mean ± SD (n = 3 from three independent batches). (**q**) Heatmap of qRT-PCR analysis showing expression of endothelial (*CD31, CD144*), pericyte (*PDGFRβ*), microglia (*IBA1*), astrocyte (*S100β, GFAP*), pluripotency (*OCT3/4, NANOG*), GTM system (*ETV2, MmPylRS*), and vasculogenesis-related genes (*MME, GLUT1*) at indicated time points.

Strikingly, we observed enriched IBA1^+^ microglia surrounding the vascular cells in 30-day vhCOs, and activated microglia showed significantly increased identified by CD11b^+^ (∼3.98%) and CD68^+^ (∼1.04%) (**Fig. 2g, h**), as well as CD31+ (∼6.26%) and CD140b+ (∼6.58%) cells (**Fig. 2h**). Additionally, to investigate the organization of the endothelium and vascular-like networks, we performed optical tissue clearing and whole-mount immunostaining, which demonstrated that CD31+ endothelial cells at day-30 vhCOs formed the vascular network compared to hCOs (**Fig. 2i, and Supplementary Fig. 4b**). By day 60, vhCOs displayed SOX2^+^/TUJ1^+^ neural cells, CD31^+^/PDGFRb^+^ vasculature, ZO1^+^ tight junctions and S100β^+^ astrocytes, which jointly constitute the BBB features (**Fig. 2j**). As expected, similar expression patterns were observed in day 90 vhCOs (**Fig. 2k**), especially more obvious vascular luminal structures (**Fig. 2l**). In day 90 vhCOs, immunostaining for GFAP and AQP4 displayed end-feet structure (**Supplementary Fig. 5a**). ECs and pericytes organized in a typical cytoarchitecture in day-90 vhCOs (**Supplementary Fig. 5b**). Moreover, CD31^+^ populations were markedly increased in vhCOs (from ∼7.7% in 60-day to ∼9.27% in 90-day) (**Fig. 2m, n**). We also observed enriched IBA1^+^ microglia from different batches surrounding the vascular cells in 90-day vhCOs, and activated microglia showed significantly increased identified by CD11b^+^ (∼4%) and CD68^+^ (∼3%) (**Fig. 2o, p and Supplementary Fig. 6a-e**). Additional microglia markers, P2RY12, TREM2 and TMEM119 were also observed in day-90 vhCOs by immunostaining (**Supplementary Fig. 6f**). To assess microglia functionality, day-90 vhCOs were treated with 100 ng/mL LPS for 24 hours. ELISA analysis of the supernatant revealed significantly increased secretion of TNF-α and IL-6 in LPS-stimulated vhCOs, indicating the functional immune responses of microglia within vhCOs (**Supplementary Fig. 6g, h**). qRT-PCR profiling showed the progressive upregulation of genes for endothelial cells (*CD31, CD144*), pericytes (*PDGFRβ*), microglia (*IBA1*), astrocytes (*S100β, GFAP*), GTM (*ETV2, MmPylRS*) and vasculogenesis (*MME, GLUT1*), along with declined expression of pluripotency genes (*OCT3/4, NANOG*) (**Fig. 2q, and Supplementary Fig. 4c**). However, day-30 cerebral organoids derived from H9^GTM^ without NAEK treatment did not exhibit CD31, IBA1, or PDGFRβ expression (**Supplementary Fig. 4d**). Immunostaining for TUJ1 and Synapsin showed a higher number of mature neurons in day-90 vhCOs compared to hCOs (**Supplementary Fig. 7**). TUNEL staining revealed that vascularization reduced cell apoptosis in 90-day vhCOs (**Supplementary Fig. 8**). These results suggest the successful generation of human cerebral organoids contained multicellular lineages including vasculature and microglia via GCE-T, which were not observed in hCOs (**Supplementary Fig. 9**). Similar results were obtained in iPS cells using GCE-T (**Supplementary Fig. 10**).

To further characterize the functions of vhCOs, we examined whether vhCOs formed tight junction barrier among interconnected ECs by trans-endothelial electrical resistance (TEER) analysis (**Fig. 3a**). TEER measurements were performed by placing microelectrode probes in three distinct regions of the organoid spheres (**Fig. 3b**). 30-, 60-, and 90-day vhCOs showed significantly higher TEER values than hCOs control (**Fig. 3c**). In order to assess the capability of vascular perfusability, day-30 vhCOs were placed in a bioreactor system and perfused with FITC-dextran under constant flow (**Fig. 3d**). Whole-mount immunostaining for CD31 revealed that FITC-dextran localized within ETV2-induced CD31+ vasculature, demonstrating the formation of a functional, perfusable vascular-like network in vhCOs (**Fig. 3e**). Moreover, vhCOs exhibited higher permeability of FITC-Dextran than hCOs, indicating the presence of abundant vasculature in vhCOs **(Supplementary Fig. 11a-c**). Microelectrode array (MEA) analysis revealed a higher mean firing rate (Hz) and network burst duration (Sec) in vhCOs compared to hCOs at the indicated time points (**Fig. 3f-i**). This suggested that vasculature might facilitate the neuronal maturation in day-90 vhCOs. To further investigate whether vhCOs form functional vasculature *in vivo*, we subcutaneously implanted vhCOs and hCOs into the hind limbs of immune-deficient mice and performed magnetic resonance imaging (MRI) on the xenografts and FITC-dextran perfusion (**Fig. 3j**). The transplanted vhCOs exhibited high contrast relative to the surrounding muscle and skin tissues, indicating the blood perfusion and vascular integration of vhCOs with host vessels (**Fig. 3k, l**). At 30 d.p.i., we identified FITC-dextran perfusion into the mice and human-specific CD31 expression, indicating the formation of functional blood vessels within vhCOs and their integration with host circulatory system *in vivo* (**Fig. 3m, n, and Supplementary Fig. 11d**).

**Figure 3.**
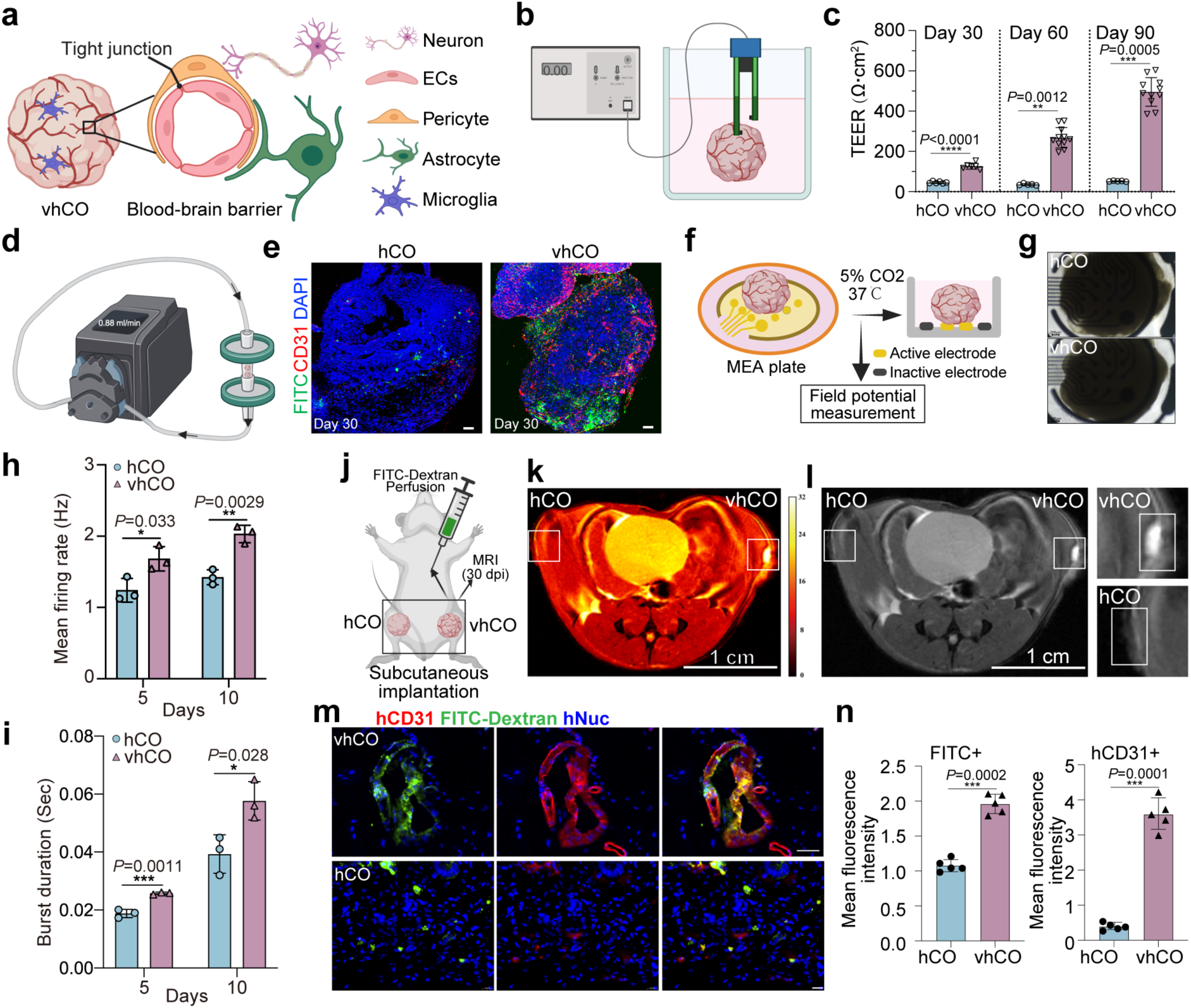
Functional characterization of vhCOs. (**a, b**) Schematic illustrations of blood-brain barrier (BBB) features and TEER measurement in vhCOs. (**c**) TEER values of hCOs and vhCOs at days 30, 60, and 90. Data are shown as mean ± SD (n ≥ 5 from three independent batches). (**d**) Schematic of FITC-dextran perfusion into organoids via a bioreactor at a flow rate of 0.88 mL/min. (**e**) Whole-mount immunostaining for CD31 in FITC-dextran-perfused vhCOs and hCOs. Representative images are shown (n = 3 from three independent batches). (**f, g**) Representative bright-field images of microelectrode array (MEA) analysis of hCOs (**f**) and vhCOs (**g**). Scale bars, 50 µm. (**h**) Weighted mean firing rate (Hz) of hCOs and vhCOs at indicated time points. Data are shown as mean ± SD (n = 3 from three independent batches). (**i**) Network burst duration (Sec) of hCOs and vhCOs at indicated time points. Data are shown as mean ± SD (n = 3 from three independent batches). (**j**) Schematic of subcutaneous implantation of hCOs and vhCOs into the right and left hindlimbs of immunodeficient mice, followed by MRI and FITC-dextran perfusion analysis at 30 days post-implantation (d.p.i.). (**k**) In vivo T2-weighted MRI of implanted control hCOs and vhCOs at 30 d.p.i. (**l**) Anatomical images of explanted hCOs and vhCOs at 30 d.p.i. Scale bar, 1 cm. Data are representative of three independent experiments. (**m, n**) Immunostaining for human-specific CD31 (red) and human nuclei (hNuclei, blue) in FITC-perfused grafts at 30 d.p.i. (**m**) and quantification of mean fluorescence intensity (**n**). The red signal in the hCO is background. Data are shown as mean ± SD (n = 5 from three independent batches). Scale bars, 100 µm. n = 3 animals and six organoids from three independent batches for MRI and FITC-perfusion analyses.

### Single-nucleus transcriptomic analysis of vhCOs

To further profile the transcriptome and cell lineage trajectories in vhCOs, we performed single-nucleus RNA sequencing (snRNA-seq) analysis of 90-day organoids (**Fig. 4a**). After quality control, a total of 17574 cells from vhCOs and 11868 cells from hCOs were retained for Uniform Manifold Approximation and Projection (UMAP) analysis to identify cell clusters. hCOs exhibited only 7 major clusters mainly from neural lineage (**Fig. 4b, c and Supplementary Fig. 12a**). In contrast, vhCOs showed 12 major clusters, including intermediate progenitor cells (IPC), radial glia (RG), glioblast (GB), neuroendocrine cells (NE), epithelial cells (Epi), neurons, endothelial cells (Endo), microglia, and other mesodermal lineages (**Fig. 4d, e and Supplementary Fig. 12b**). It indicated that vhCOs contained a greater diversity of cell types, especially the endothelial cells and microglia (**Supplementary Fig. 12c**). Interestingly, the mesodermal-hematopoietic progenitors (Meso-hemato) expressing KDR and GATA4 were only detectable in vhCOs. It indicates that the Meso-hemato cell cluster may give rise to hematopoietic cells and endothelium that support vascular development^[21–26]^, as well as to tissue resident macrophages that are essential for tissue homeostasis^[27,28]^. In addition, analysis of genes differentially expressed in vhCOs showed high level of GATA4, as an endothelial/hematopoietic transcription factor, which is important for the hemogenic competence of mesodermal progenitors (**Fig. 4e**). hCOs differentiation follows a trajectory from progenitor cells to neurons (**Fig. 4f, h, i**), while the differentiation trajectory of vhCOs distinctly separates ectodermal and mesodermal lineages (**Fig. 4g, j, k**). This analysis revealed the coordinated differentiation of both vascular progenitors (meso-hemato) and vascular-supporting mesodermal lineages (perivascular). Notably, vhCOs initiated vascularization at earlier developmental stage, while exhibited high expression of neuronal maturation genes at late stage (**Fig. 4i, k**). Moreover, microglia emerged at earlier stage, and mesenchymal/extracellular matrix (ECM) genes linked to vascularization were highly upregulated during mid-development (**Fig. 4k**). This finding suggests that paracrine signaling from endothelial cells may be essential for maintaining microglia during organoid development. These results are further supported by the RNA velocity analysis (**Fig. 4f, g**). Additionally, a cluster-cluster communication network map revealed that vhCOs displayed more extensive cell-cell interactions compared to hCOs (**Fig. 4l, m**). Comparison of communication patterns between hCOs and vhCOs showed that vhCOs not only significantly activated vascularization-related pathways (e.g., JAM2, BMP, PTN), but also demonstrated a more enriched communication network than that in hCOs in neural development (e.g., SEMA, EPHA), tissue development (e.g., NOTCH, FGF, IGF), and cell migration (e.g., FN1, SPP1, CD99, TENASCIN) (**Supplementary Fig. 12d**). The most pronounced differences were observed in ECM-related pathways (COLLAGEN, LAMININ) and neural development pathways (NRG, NRXN, NCAM) (**Supplementary Fig. 12e**). In addition, we analyzed the expression levels and cellular distribution of *ETV2, TREM2, P2RY12, TMEM119*, and *IBA1 (AIF1)* from snRNA-seq (**Supplementary Fig. 13a, b**). RUNX1, a key regulator of microglial development, is expressed exclusively in mesodermal (perivascular) lineages and microglia, reflecting the definitive identity and endogenous origin of these cells (**Supplementary Fig. 13c**). Quantitative RT-PCR validated the expression of *TREM2, P2RY12 and TMEM119* (**Supplementary Fig. 13d**). These pathways mediated interactions between neural, immune and vascular cells, which may contribute to neurogenesis and organoid maturation. These findings validated that GCE-T enabled the generation of cerebral organoids with temporal coordination of multiple cell lineages including vasculature and microglia.

**Figure 4.**
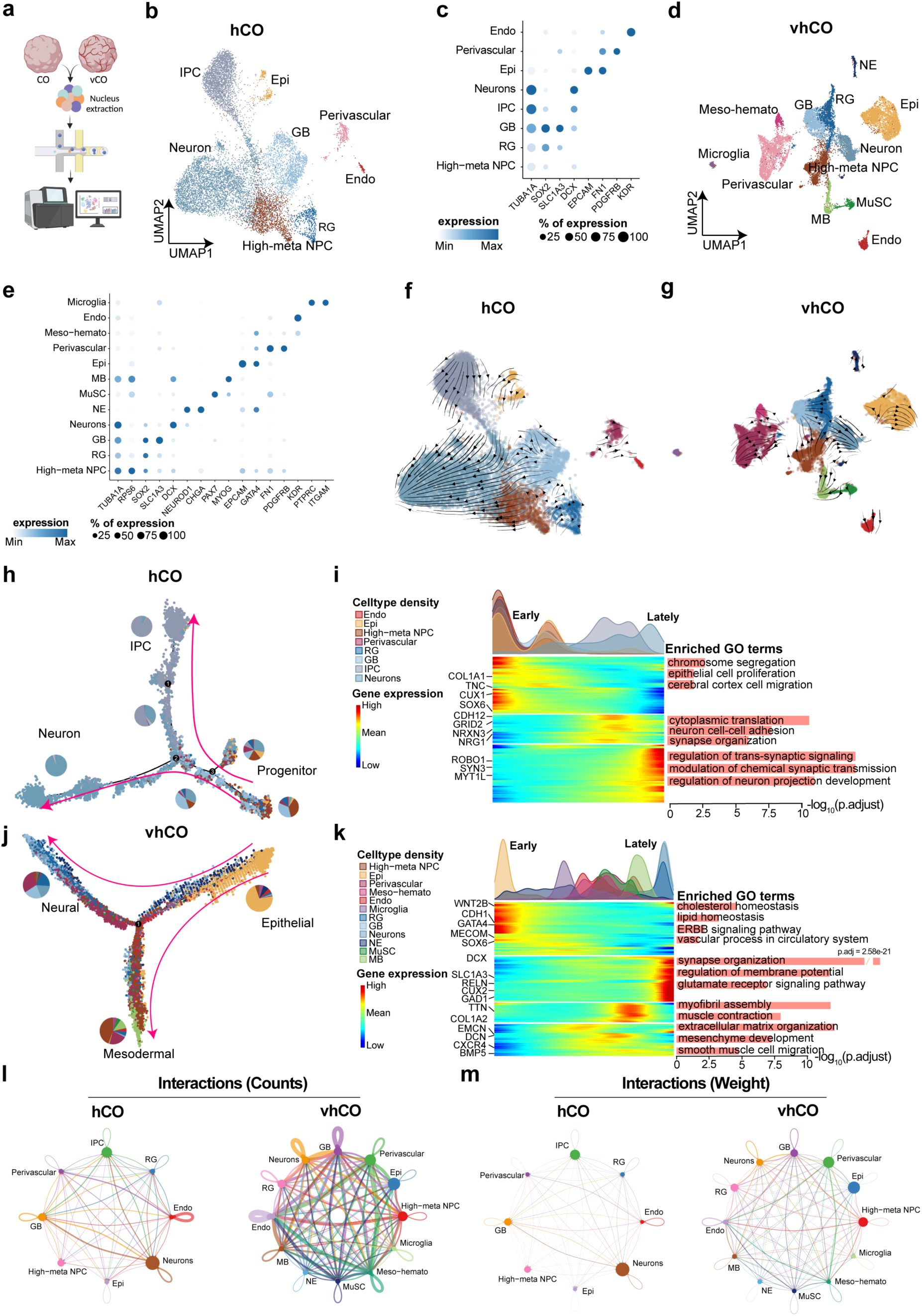
Single-nucleus transcriptomic characterization of hCO and vhCO at day 90. (**a**) Schematic of the snRNA-seq workflow. (**b**) UMAP plot showing major cell types identified in hCOs. (**c**) Marker genes defining cell types in hCOs. (**d**) UMAP plot showing major cell types identified in vhCOs. (**e**) Marker genes defining cell types in vhCOs. Abbreviations: IPC, intermediate progenitor cells; GB, glioblasts; RG, radial glia; Endo, endothelial cells; Epi, epithelial cells; NPC, neural progenitor cells; NE, neuroendocrine cells; MB, myoblasts; MuSC, muscle satellite cells. (**f**) RNA velocity-based differentiation stream for hCOs, revealing a robust neuronal differentiation trajectory. (**g**) RNA velocity-based differentiation manifold for vhCOs, illustrating differentiation trajectories including RG to GB and neurons, MuSC to MB, and progenitors to perivascular cells, endothelial cells, or microglia. (**h**) Pseudotime trajectory analysis showing differentiation from neural progenitors to neurons in hCOs. (**i**) (Top) Density distribution of cell types along the pseudo time trajectory. (Bottom) Dynamic gene expression along the trajectory and corresponding GO enrichment results for each gene module. (**j**) Pseudotime trajectory analysis showing bifurcation of progenitors (epithelial) into neural (ectodermal) and mesodermal lineages in vhCOs. (**k**) (Top) Density distribution of cell types along the pseudotime trajectory, with microglia and vasculature-associated mesodermal cells emerging at early time points. (Bottom) Dynamic gene expression along the trajectory and GO enrichment results for each module. (**l, m**) Cell-cell communication networks in hCOs (**l**) and vhCOs (**m**). Plot parameters were standardized across conditions to ensure comparability.

### vhCOs possess multi-brain-regional identity with diverse neuronal subtypes

To examine whether these phenotypic changes align closely with human brain development, we applied deep generative modeling and transfer learning to map our data onto a human fetal whole-brain atlas^[29]^ (see Methods). This analysis showed that neural lineages in vhCOs exhibited broader regional distribution, including the cerebellum, midbrain, and brainstem, whereas hCOs were largely confined to the cortex/forebrain (**Supplementary Fig. 14a**). To validate this finding at the gene expression level, we defined brain-region-specific gene sets from the fetal brain atlas using both a lenient method (one-vs-all comparison) and a stringent method (pairwise comparisons with intersecting results) (**Supplementary Fig. 14b**). Gene Set Variation Analysis (GSVA) was then applied to assess enrichment within the neural lineages of hCOs and vhCOs. Both sets confirmed that vhCO neural lineages exhibit multi-regional identity (**Fig. 5a and Supplementary Fig. 14c**).

**Figure 5.**
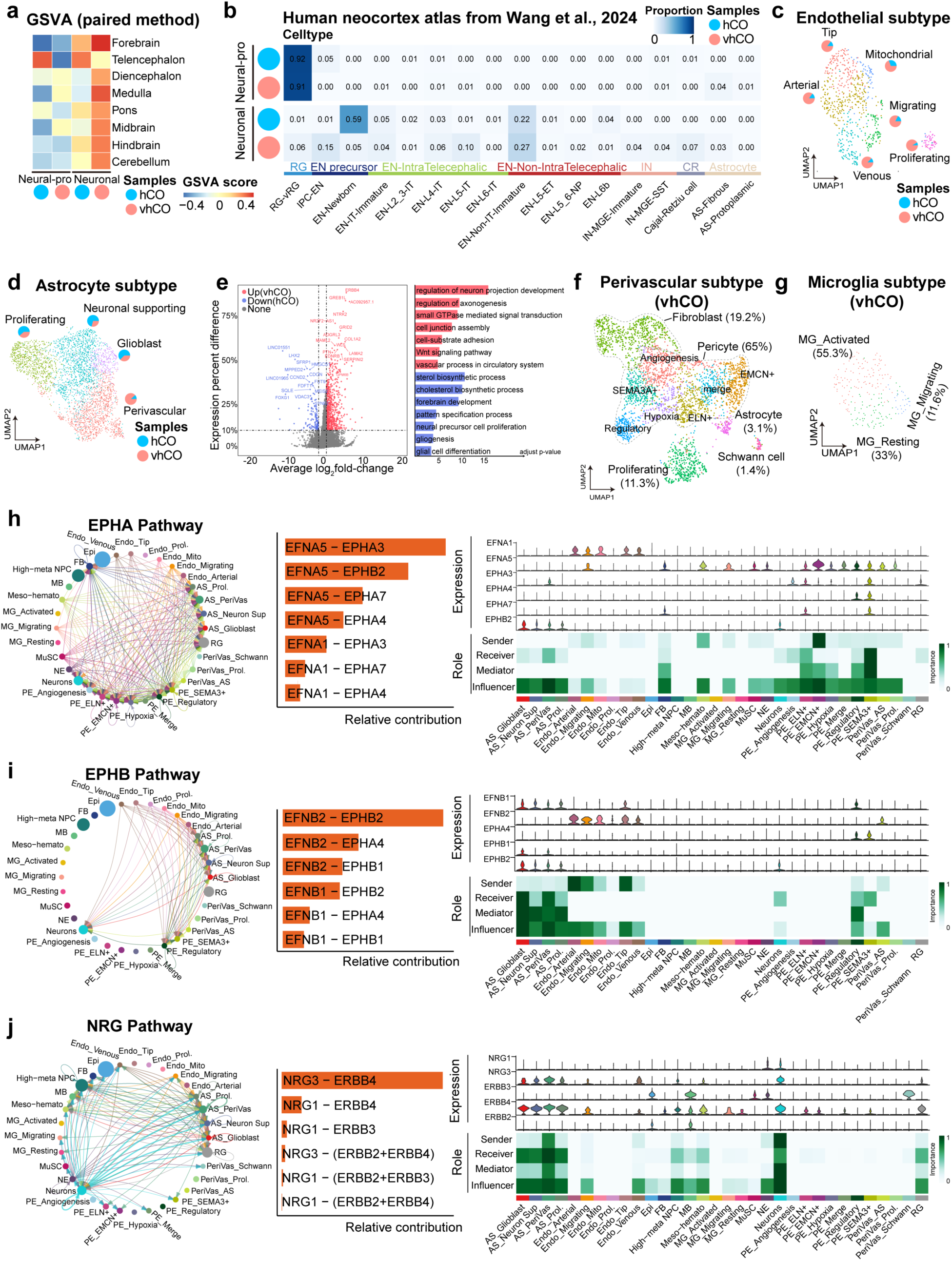
vhCOs exhibit diverse brain-region and neuronal characteristics with enhanced neurovascular interactions compared to hCOs. (**a**) Gene Set Variation Analysis (GSVA) enrichment scores for brain-region-specific gene sets (**see Fig. S14b**) in the neural lineages of hCOs and vhCOs. (**b**) Mapping of hCO and vhCO datasets onto a human neocortex reference atlas using a deep generative model with transfer learning (**see Methods**). Heatmap shows the proportion of cells from each dataset mapped to reference cell types. (**c**) Subcluster classification of glioblast (GB, astrocyte-like) populations in hCOs and vhCOs. Pie charts show the proportion of cells from each condition within each subcluster. (**d**) Differential gene expression and GO enrichment analysis for GB populations between hCOs and vhCOs. The x-axis represents log-transformed fold change; the y-axis indicates the difference in expression proportion between conditions. (**e**) Subcluster classification of endothelial (Endo) populations in hCOs and vhCOs. Pie charts show condition proportions within each subcluster. (**f**) Subcluster classification of perivascular cell populations in vhCOs. (**g**) Microglia subtypes identified in vhCOs. (**h-j**) Cell-cell communication networks within vhCOs highlighting key neurovascular interaction pathways: EPHA (**h**), EPHB (**i**), and NRG (**j**). For each pathway: (Left) Chord diagrams illustrating intercellular communication; (Middle) Signal intensity contribution of each ligand-receptor pair; (Right) Functional roles of each cell type (Bottom) and corresponding gene expression (Top).

To determine the neural lineage specification underlying multi-brain-regional identity, we mapped our data onto a human neocortical atlas^[30]^. vhCOs exhibited broader neuronal subtype diversity, comprising substantial proportions of mature excitatory neurons (52%), inhibitory neurons (8%), and Cajal-Retzius (CR) cells (7%). In contrast, hCOs were predominantly composed of excitatory neuron precursors (60%), with marked deficiencies in both inhibitory neurons and CR cells (**Fig. 5b**). To assess maturity of organoids, we mapped the entire hCO and vhCO datasets onto a post-conception week 5 human gastrulation atlas^[31]^. This analysis revealed higher presence scores for hCOs during gastrulation (particularly in neural crest), indicating greater maturation in vhCOs compared to hCOs (**Supplementary Fig. 14d**). Collectively, these results demonstrate that vhCOs possess enhanced multi-brain-regional identity, increased neuronal subtype diversity, and advanced maturation relative to hCOs.

### vhCOs exhibit enhanced neurovascular crosstalk

*In vivo*, neurovascular crosstalk is responsible for maintaining local functionality and homeostasis in the brain. We further analyzed intercellular communication networks in vhCOs and hCOs. Significantly, vhCOs exhibited enhanced signaling between EC, GB, neurons and perivascular cells compared to hCOs (**Supplementary Fig. 15a**). Subcluster analysis confirmed that GBs in both organoid types were predominantly astrocytes (**Fig. 5d and Supplementary Fig. 14f**). Critically, hCO astrocytes aligned primarily with neuro-supportive subtypes, while vhCO astrocytes exhibited a perivascular-associated transcriptional signature (**Fig. 5d**). Differential gene expression (DEG) analysis and GO enrichment of the GB populations in hCOs and vhCOs revealed distinct functional roles. hCO astrocytes were enriched for small-molecule metabolism, whereas vhCO astrocytes were associated with neuronal projection, axonogenesis, and vascular development (**Fig. 5e**). When comparing endothelial subpopulations, vhCOs displayed pronounced expansion of EC population with distinct arterial-, tip-, and venous-type ECs (**Fig. 5c and Supplementary Fig. 14e**). Perivascular populations in vhCOs comprised multiple pericyte subtypes, fibroblasts, astrocytes, and Schwann cells (**Fig. 5f and Supplementary Fig. 14g**). Additionally, we identified microglia in vhCOs comprising three distinct subtypes: activated (55.3%), resting (33%), and migrating (11.6%) cells (**Fig. 5g and Supplementary Fig. 14h**). Notably, Microglia-vascular interactions were characterized by activated microglia engaging via SPP1-integrin binding and ICAM signaling, which may modulate vascular inflammatory/growth. Migrating microglia secrete POSTN (periostin), which may stabilize vasculature through synergistic interactions with ECM-related signaling (**Supplementary Fig. 17**).

To investigate the roles of various subclusters in neurovascular interactions and BBB formation, we constructed a subpopulation-level cell communication network for vhCOs (**Supplementary Fig. 15b, c**). Non-negative matrix factorization (NMF) identified five output modules and four input modules, revealing enhanced neurovascular signaling across multiple subpopulations (**Supplementary Fig. 15d, f**). Specifically, pericytes and ECs served as primary signal senders, regulating neurovascular development through EPHA/EPHB signaling (**Fig. 5h, i**). The NRG pathway was predominantly mediated by astrocytes, neurons and ECs (**Fig. 5j**). Different pericyte subtypes played distinct roles in neurovascular signaling: hypoxia-responsive pericytes promoted angiogenesis via ANGPTL signaling, likely localized near lumen-deficient vascular sprouts (**Supplementary Fig. 16a**). The JAM pathway, primarily involving EMCN+ pericytes, astrocytes, and ECs, suggested functional BBB maturation (**Supplementary Fig. 16a**). Additionally, we identified other distinct signaling networks, including COLLAGEN, LAMININ, NCAM, PTN, and NOTCH pathways, mediating cellular interactions, later-stage developmental and migratory processes in vhCOs (**Supplementary Fig. 16a**).

Furthermore, we investigated whether the endothelial subclusters influences the development of cerebral organoids. We identified extensive signaling events within endothelial subclusters and constructed dedicated endothelial-endothelial interaction network (**Supplementary Fig. 15c**). It revealed that Tip and Arterial ECs drive angiogenesis and dominate intravascular crosstalk through NOTCH (DLL4, DLK1), PTPRM, and PGF-FLT1 pathways, suggesting arterial-type ECs as primary mediators of neovascularization in vhCOs (**Supplementary Fig. 16b**). Notably, two astrocyte subclusters demonstrated BBB-related specialization: PeriVas_AS in the perivascular cluster and AS_PeriVas in the GB cluster (**Fig. 5d, f**). Interestingly, DEGs and signal pathways of PeriVas_AS were enriched in ECM organization and vascular/mesenchymal development, while AS_PeriVas showed greater involvement in neural developmental processes (**Supplementary Fig. 16c, d**). It suggested that PeriVas_AS may directly contribute to BBB structural integrity, while AS_PeriVas facilitates BBB-neural system crosstalk.

In order to validate the microglia-vascular crosstalk, day 90 vhCOs were treated with A205804, an E-selectin/ICAM-1 inhibitor. This inhibitor did not affect TEER values and FITC permeability in vhCOs, but it reduced the adhesion of microglia to vasculature and microglia activation (**Supplementary Fig. 18a-e**). The results revealed the ICAM-integrin signaling might be functionally necessary for the microglia-vascular crosstalk and coordinated development in vhCOs.

### vhCOs show neurovascular dysfunction and impaired microglia development to ZIKA virus infection

ZIKV is a known cause of microcephaly and neurological disorders. Previous studies have employed brain organoid to study ZIKV infection and probe the pathogenesis of virus-induced microcephaly^[32–35]^. However, these models are inherently limited by the absence of functional vascular networks and immune components, which are critical modulators of ZIKV neuroinvasion and host responses *in vivo*. To gain new insights using the vhCOs model, we infected 90-day vhCOs with ZIKV for 24 h, followed by 6 days of observation (**Fig. 6a**). This late-stage infection was chosen to align with snRNA-seq data and maximize cellular diversity. Infected organoids showed significant reduction in size versus control (**Fig. 6b, and Supplementary Fig. 19a**). Immunostaining revealed ZIKV-induced disruption, including damaged endothelial network (**Fig. 6c, d**), decreased SOX2+ neural progenitor cells (NPCs) (**Fig. 6c**), preferential infection of NPCs and TUJ1^+^ neurons (**Fig. 6c, e**), and impaired microglia development (**Fig. 6f, and Supplementary Fig. 19b**). Day-90 hCOs were infected with ZIKV or mock virus. ZIKV preferentially infected neural progenitor cells (NPCs) and TUJ1+ neurons, leading to increased cell death, reduced proliferation, and decreased neuronal cell-layer volume. This is consistent with previous report^32^. Of note, we did not observe any effects of ZIKV on endothelial cells or microglia in hCOs, as these cell types are absent in this model (**Supplementary Fig. 19c-e**). Sn-RNAseq of infected vhCOs further demonstrated the reduced cell-type diversity, diminished perivascular/mesenchymal characteristics, and impaired microglia development (**Fig. 6g**). Transcriptional analysis revealed activation of immune/inflammatory factors (STAT3, NFKB2, ATF4), indicating cellular stress (**Fig. 6h**). Neurodevelopmental pathways (NCAM, NRXN, NRG) were severely downregulated (**Fig. 6i**). Mapping analysis showed decreased neuronal identity scores and increased radial glia (RG) abundance (**Supplementary Fig. 20a**). Viral genome localization (86.3% cell debris and 9.7% intact neural cells; **Supplementary Fig. 20b**) confirmed ZIKV’s neurotropism drives cell death and developmental arrest. Comparative analysis with fetal brain revealed that 56% of ECs in infected vhCOs acquired immune-lineage transcriptional signatures (**Fig. 6j**). Viral-response genes were enriched in NPCs, Mural cells, ECs, Fibroblast, and MSCs (**Fig. 6k**). Notably, canonical innate immune pathways (e.g., JAK, NF-κB, IFN-α, IFN-γ) were enriched in ECs, while cellular stress pathways (e.g., apoptosis, autophagy, neurodegeneration and viral replication) were enriched in NPCs, neurons and RG (**Fig. 6k**). Subcluster analysis identified transcriptional profiles of immune hyperactivation in ZIKV-infected ECs (**Fig. 6m**). These findings suggest that ZIKV infection can increase innate immunity in ECs and propagate anti-viral response, which may regulate endothelial dysfunction.

**Figure 6.**
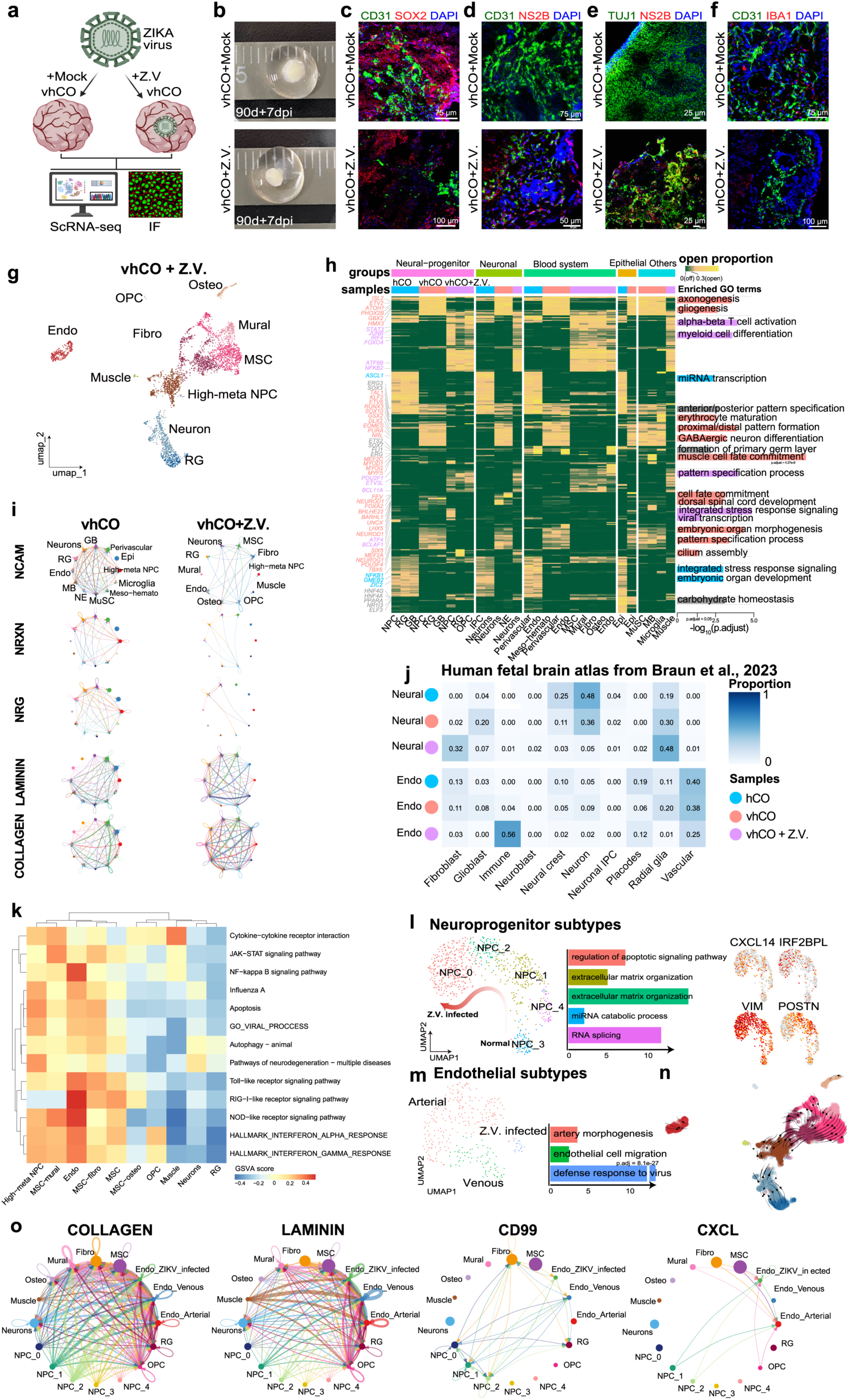
Zika virus infection disrupts neural development, damages microglia, activates vascular immunity, and induces mesenchymal transition in vhCOs. (**a**) Schematic workflow of Zika virus (ZIKV) infection in hCOs and vhCOs and subsequent characterization. (**b**) Bright-field images showing size changes of hCOs and vhCOs at 7 days post ZIKV infection (d.p.i.). (**c-f**) Immunostaining of CD31+/SOX2+ (**c**), CD31+/NS2B+ (**d**), TUJ1+/NS2B+ (**e**), and CD31+/IBA1+ (**f**) cells at 7 d.p.i. Scale bars, 75 µm, 100 µm, 50 µm, 25 µm, respectively. (**g**) UMAP visualization of major cell types in ZIKV-infected vhCOs (vhCO+ZIKV). (**h**) Transcription factor (TF) activation scores across cell types in hCOs, vhCOs, and vhCO+ZIKV. Clustering was performed using Euclidean distance, with GO enrichment for TFs in each module (right). (**i**) Cell-cell communication networks of key neural developmental signaling pathways across major cell types in vhCOs and vhCO+ZIKV. (**j**) Mapping of neural lineage and endothelial populations from hCOs, vhCOs, and vhCO+ZIKV onto a human fetal brain atlas, revealing mesenchymal transition of neural lineages and endothelial immune activation in vhCO+ZIKV. (**k**) Enrichment of viral infection-associated signaling pathway genes (from KEGG and MSigDB) across cell types in vhCO+ZIKV. (**l**) Subcluster classification of neural progenitor cells (NPCs) in vhCO+ZIKV (left), GO enrichment of corresponding marker genes (middle), and expression of mesenchymal (VIM, POSTN) and immune-related (CXCL14, IRF2BPL) genes (right), revealing a normal-to-mesenchymal-to-viral infection trajectory. (**m**) Subcluster classification of endothelial cells in vhCO+ZIKV (left) and GO enrichment of corresponding marker genes (right), identifying ZIKV-infected endothelial subpopulations. (**n**) RNA velocity-based differentiation stream plot showing abnormal neurodevelopmental trajectories, including NPCs shifting toward a mesenchymal lineage and multi-source fibroblast-like mesenchymal cell differentiation. (**o**) Subcluster-level cell-cell communication in vhCO+ZIKV. Mesenchymal-like and ZIKV-infected NPCs (NPC_0,1,2)—but not normal NPCs (NPC_3,4)—predominantly signal through extracellular matrix components (COLLAGEN, LAMININ) and cell migration/immune pathways (CD99, CXCL).

Notably, a high proportion of NPCs (32%) in ZIKV-infected vhCOs acquired transcriptional profiles converging with fetal brain fibroblasts (**Fig. 6j**). UMAP visualization positioned these NPCs closer to mesenchymal rather than neural lineages (**Fig. 4b, c and Fig. 6g**). Spearman correlation also confirmed significantly reduced association between NPCs and neural clusters (**Supplementary Fig. 20d**). Subcluster analysis defined a distinct NPC differentiation trajectory: from normal metabolic state (NPC_3,4) to mesenchymal development (NPC_1,2) in ZIKV-infected vhCOs (**Fig. 6l**). The NPC-mesenchymal transition progress was validated by GO enrichment, typical gene expression patterns, gene set enrichment analysis and RNA velocity-based differentiation trajectories (**Fig. 6l, n and Supplementary Fig. 20e**). In transitional mesenchymal cells, significantly enriched genes were implicated in ECM organization and cell migration pathways (**Fig. 6o and Supplementary Fig. 20h, i, j, k and 21**). Additionally, the development of NPCs and neurons was severely disrupted after ZIKV infection (**Supplementary Fig. 20f, g**). Overall, these findings suggest that ZIKV could cause neurovascular dysfunction, NPC-mesenchymal transition and impaired microglia development, which may provide insights into ZIKV infection induced neurodevelopmental disorders.

## Discussion

This work describes the first use of site-specific protein engineering to regulate ETV2 expression for the generation of vascularized brain organoids. Our system enables precise translational control, exhibiting minimal leakage and rapid responsiveness. By temporal control of ETV2 expression at the translational level, vhCOs achieved coordinated differentiation of both vasculature and microglia-like cells as well as functional BBB features, faithfully recapitulating human fetal brain development. Moreover, vhCOs exhibited a functional connection to the host circulatory system, forming perfusable vascular networks *in vivo*. SnRNA-seq analysis revealed enhanced neurovascular interactions and multi-brain-regional identities in vhCOs. Specially, the microglia-like cells comprised three distinct subtypes and synergistically modulated vascular development via microglia-vascular interactions. This strategy provides a powerful platform to engineer brain organoids, facilitating the study of neurodevelopment and pathophysiology of the fetal brain.

Researchers have produced brain organoids integrated with vascular or immune cells using various approaches^[10–13]^. For instance, Sun et al. fused separately grown vessel and brain organoids to achieve functional BBB-like structures and microglia incorporation^[12]^. However, differences in germ layer origins may limit fusion efficiency, and the multi-step process is resource-intensive. Abud et al. reported a co-culture method in which pre-differentiated microglia or hematopoietic progenitors were co-cultured with cerebral organoids. This is an excellent method for introducing mature microglial function but models an “invasion” or “engraftment” scenario^[36]^. Ormel et al. reported a co-differentiation method to innately generate microglia within a cerebral organoid by subtle modifications in previous protocol (e.g., differentiation path, heparin dosage, matrix embedding time, and media composition)^[37]^. The internally generated functional microglia-like cells produced by GCE-T present a distinct and complementary approach through translation regulation of ETV2, which acts as a pioneer factor for a mesodermal-hematopoietic progenitor state. Furthermore, microglia within vhCOs survived robustly in standard organoid maintenance media without any added trophic factors. This aligns with previous reports demonstrating that endogenous cell populations within the organoid, such as excitatory neurons expressing IL34, radial glia expressing CSF1, and dividing progenitors expressing TGFβ, provide the necessary factors for microglial survival^[38]^. By introducing PTCs into surface-exposed residues of *ETV2*, we achieved precisely controllable and rapid expression of full-length ETV2 in the engineered hPSCs and their derived brain organoids. Different from previous approaches^[13]^, the GCE-T-based method enables the coordinated development of multiple cell lineages in organoids, including vasculature and microglia-like cells, representing a more valuable model system.

snRNA-seq analysis revealed a relative decline in neural lineage proportions in vhCOs, which can be attributed to mesodermal population expansion rather than neurodevelopmental deficits. As shown in Fig. 5d, vhCOs exhibit a broader range of neuronal subtypes, including excitatory neurons, inhibitory neurons, and Cajal-Retzius cells, compared to hCOs. Mapping to a human gastrulation atlas shows vhCOs are more developmentally advanced than hCOs (Supplementary Fig. 14d). MEA analysis revealed higher firing rates and network burst duration in vhCOs, indicating that vascularization supports rather than hinders neuronal functionality (Fig. 3h, i,). Furthermore, analysis of cell-cell communication (Fig. 5j-k and Supplementary Fig. 15) reveals enhanced signaling interactions between vascular, glial, and neuronal populations in vhCOs, indicating that mesodermal expansion promotes the establishment of a more physiologically relevant microenvironment. Collectively, these multi-dimensional data demonstrate that while the introduction of mesodermal lineages alters cellular composition, the fidelity, diversity, and functionality of neuronal networks are not only preserved but are significantly enhanced in vhCOs.

This interpretation is supported by the absence of delayed neural maturation, loss of neuronal identity, or signs of cellular stress. Notably, this system promotes progressive mesodermal specification (potentially originating from GATA4+ epithelial progenitors) rather than direct neural-to-vascular transdifferentiation. The expanded mesodermal population exhibited extensive crosstalk with neural lineages, including activated neurovascular interactions associated with ECM remodeling, neurodevelopment, and cellular growth, collectively fostering maturation and functionality of neural networks. Furthermore, we found that GCE-T-mediated translational regulation of ETV2 activated NOTCH and BMP signaling, which induced the differentiation of mesoderm/hematopoietic progenitor cells that subsequently give rise to both ECs and microglia. To assess physiological relevance, we compared vhCOs with fetal brain tissue, demonstrating robust multi-regional identity and enhanced neuronal maturation. These findings demonstrate that GCE-T enables efficient generation of vascular, immune, and mesodermal cells without compromising neuronal development.

In this study, ZIKV infection led to damaged vasculature, disrupted neurogenesis, abnormal microglia development, and activation of innate immune signaling in vhCOs, revealing the neurovascular dysfunction in the fetal brain after ZIKV exposure. It elucidates unexplored pathogenesis of viral pathogenesis in the human fetal brain involving neurovascular interactions and immune compartments. Notably, when microglial development was markedly delayed, ECs were the primary virus-resistant cell population, and infection spread from neural lineages to the vascular cells. Furthermore, we found that ZIKV induced mesenchymal transition in NPCs, characterized by fibroblast-like morphology, and this transition may represent a survival adaptation in the chronically ZIKV-infected fetal brain. These new findings revealed multicellular interactions mediated pathogenesis of ZIKV-induced neurodevelopmental disorders.

Taken together, this GCE-based methodology provided a new paradigm to generate brain organoids with functional vasculature and immune components through site-specific ETV2 protein engineering, which faithfully recapitulate developmental trajectory of the human fetal brain. The novelty of this work lies in its ability to generate cerebral organoids with coordinated development of multiple cell lineages including vasculature and microglia using GCE-T, which is not readily achieved by existing methods. This work provides a novel synergistic strategy to engineer high-fidelity brain organoids, advancing organoid research and biomedical applications.

## Materials and Methods

### Cell lines and culture

H9 hESCs (WiCell, no. WB68075) were purchased from WiCell Research Institute. iPSCs from skin fibroblast were purchased from Coriell Institute (GM25256). hPSCs (H9s and iPSCs) were maintained in a six-well plate coated with Matrigel (354277, Corning) in mTeSR1 (STEMCELL) medium. Cells were passaged every 4 days with ReLeSR (STEMCELL). The medium was changed daily. Cells were routinely checked for the expression of pluripotency markers, OCT3/4 and NANOG, their capability to form teratomas in immunodeficient mice, their karyotypes, bacterial and mycoplasma contaminations.

### Genetic translation machinery (GTM) plasmid construction and transfection

The PiggyBac transposon vector (SBI) was used for insertion of gene expression cassettes. The expression cassettes encoding aminoacyl-tRNA synthetases and its bio-orthogonal tRNAs were constructed as follows: PiggyBac-MmPylRS-(7sk-tRNA^Pyl^)×4-Puro and incorporated into PiggyBac transposon vector by Gibson Assembly method, respectively, followed by site-directed mutations of the tRNA anticodon to CUA. We then generated PiggyBac-ETV2 containing TAG sites at 122aa by site-directed mutations method. The unnatural amino acid, NAEK, was synthesized as previously reported^[39]^. PiggyBac-MmPylRS-(7sk-tRNA^Pyl^)×4-Puro and PiggyBac-ETV2*-Neo were co-transfected into H9 or iPS cells by electroporation (Neon NxT) following parameters of 1050 V, 10 ms and 3 pulses to construct stable H9^GTM^ or iPSC^GTM^ line. For NAEK induction assay, the medium was changed to fresh mTeSR1 with or without 1 mM NAEK. Cells were grown for an additional 48 h and the qPCR or western blot was used to detect ETV2, MmPylRS and tRNA^Pyl^ expression.

### Generation and culture of cerebral organoids

Cerebral organoids were generated from hESCs according to previous protocols with minor modifications. Briefly, for embryoid body (EB) formation, hESCs were washed twice with DPBS, incubated with Accutase for 5 minutes, and dissociated into single cells. 3000 single cells were seeded in each well of low attachment 96-well U-bottom plate in mTeSR1 medium containing 10 µM ROCK inhibitor and centrifuged at 100 g for 3 min, then medium was half changed every other day. On day 4, EBs were transferred to low attachment 24-well plate in neural induction medium containing DMEM-F12 (GIBCO) with 1% N2 supplement (GIBCO), 1% Glutamax supplement (GIBCO), 1% MEM-NEAA (GIBCO) and 1 µg/mL Heparin (GIBCO), and medium was changed after 48 h. On day 8, EBs were transferred into Matrigel (356234, Corning) droplets as previously described and cultured in cerebral organoid differentiation media containing 50% DMEM-F12, 50% Neurobasal, 200× N2 supplement, 100× B27 supplement without Vitamin A, 0.025% Insulin (GIBCO), 100× Glutamax-I supplement, 200× MEM-NEAA, 100× penicillin-streptomycin, and 0.035% 2-Mercaptoethanol, and medium was changed after 48 h. On day 12, organoids were transferred to orbital shaker (Corning) in cerebral organoid differentiation media with Vitamin A, medium was changed every 4 days.

### Generation of vascularized human cerebral organoids (vhCOs)

Based on the strategy of cerebral organoid generation mentioned above, the starting cells were H9^GTM^ or iPSC^GTM^. On day 18, 1 mM NAEK was added into cerebral organoid differentiation media for 10 days, then maintaining NAEK at a low concentration level (0.1 mM) in the medium and continued culturing until analysis.

### Microelectrode arrays (MEA)

Add 100 µL of DMEM/F12 containing 1% Matrigel (Corning, 354277) to the central detection well of the MEA plate, and coat in a 37 °C CO₂ incubator for at least 1 hour. Day 90 brain organoids were first treated with Cell Recovery Solution (354253, Corning) for 15 min at 4 °C to remove the Matrigel. Carefully aspirate the coating solution and transfer the organoids to the central detection well of the MEA plate using a sterile Pasteur pipette. Remove excess culture medium, then overlay each organoid with 15-20 µL of 100% Matrigel using a pipette. Gently push the organoids to the central detection area of the MEA plate with the pipette tip, ensuring they are properly positioned in the center. Transfer the plate to a 37 °C CO₂ incubator and allow the Matrigel to solidify completely for 30 minutes. After the organoids have adhered, slowly add 200 µL of culture medium containing 50% DMEM-F12, 50% Neurobasal, 200× N2 supplement, 0.025% Insulin (GIBCO), 100× Glutamax-I supplement, 200x MEM-NEAA, 100x penicillin/streptomycin, 0.035% 2-Mercaptoethanol and 100× B27 supplement with Retinoic Acid (RA), to fully cover them, and incubate overnight in a 37°C CO₂ incubator. The next day, confirm under a microscope that the organoids have adhered adequately, then supplement the medium to a total volume of 1 mL and continue culturing. Change the medium every 2–3 days. MEA recordings were performed at indicated time point at 37 °C in a Maestro MEA system with AxIS software using a bandwidth with a filter for 10Hz to 2.5 kHz cutoff frequencies. The phase contrast images of organoids seeded in the MEA plates were taken after MEA recording. The weighted mean firing rate (Hz) and network burst duration (Sec) were analyzed for each group (n=3 biological replicates).

### Embryonic body (EB) differentiation

hPSCs were suspended in DMEM+20% FBS+10 µM β-mercaptoethanol in low adhesion plate for 6 days. The cell masses were then transferred into plates coated with 0.1% gelatin and cultured in the same medium for another 6 days, followed by fixation and staining for NESTIN^+^ ectodermal, Brachyury^+^ mesodermal and FOXA2^+^ endodermal cells.

### Teratoma formation *in vivo*

The animal experiments were carried out following the protocols approved by the University of Science and Technology of China Animal Care and Use Committee. 1.0×10^6^ hPSCs were suspended in 25 µL PBS+25 µL Matrigel and injected subcutaneously at the back of the neck of the NOD-SCID mice (female, age 7 weeks). Teratoma was harvested when its size reached 2 cm. Teratoma was fixed with 4% Paraformaldehyde (PFA) for 48h, dehydrated with 70%, 95% and 100% ethanol sequentially, and de-fated with xylene for 2 h before embedded in paraffin. 10 µm thick section was cut and stained with hematoxylin and eosin for neural rosette, muscle and gut-like structure. Three mice (two teratomas per mouse) were used for teratoma assay for each hPSC line.

### Immunofluorescence staining

For cell immunofluorescence staining, the cells were fixed with 4% PFA at room temperature for 10 min, permeated with PBST (PBS with 0.1% Triton X-100) for 20 min and blocked with 1% BSA for 30 min. Then the cells were incubated with primary antibodies listed in **Table S1** at 4 °C overnight. The cells were subsequently incubated with secondary antibodies listed in Table S1 at room temperature for 1 h. The cells were mounted with mounting fluid containing DAPI (Yeason, 36308ES11). For organoid immunofluorescence staining, the organoids were fixed with 4% PFA at room temperature for 2 h, then immersed in 30% (w/v) sucrose until submersion before embedding and freezing in the Optimal Cutting Temperature (OCT) compound (Tissue-Tek). Serial 20 µm sections were obtained by cryo-sectioning of the embedded organoid at -20 °C using a cryostat (Leica). Cryosections were permeated with in PBST at temperature for 30 min and blocked with sheep serum for 1 h. The sections were incubated with primary antibodies listed in **Table S1** diluted in blocking buffer at 4 °C overnight. The slides were subsequently incubated with secondary antibody at room temperature for 1 h. The slides were mounted with mounting fluid containing DAPI. Stained sections were photographed under a LEICA STELLARIS 5 microscope. Apoptotic cells were labelled with TUNEL detection kit (C1090, Beyotime) according to manufacture instructions.

### Optical Clearing and Immunostaining of vhCOs

vhCOs were harvested and washed in 1x PBS to remove culture medium, then fixed in 4% PFA overnight at 4 °C with gentle shaking. Following fixation, organoids were washed three times in PBS for 1 hour each at room temperature to remove residual PFA. Fixed organoids were first dehydrated through a graded methanol series (20%, 40%, 60%, 80%, 100% in ddH₂O) for 1 hour per step. Samples were then bleached in a solution of 5% H₂O₂ in methanol overnight at 4 °C to reduce autofluorescence. After bleaching, organoids were rehydrated back to PBS via a descending methanol series. Permeabilization was performed by incubating the samples in PBS containing 0.2% Triton X-100 (PBST) for 2 hours at room temperature. Organoids were then blocked in PBST supplemented with 10% donkey serum and 6% bovine serum albumin (BSA) for 24 hours at 37 °C to prevent non-specific antibody binding. Samples were incubated with primary antibodies (e.g., anti-CD31) diluted in PBST with 5% DMSO and 3% donkey serum for 3-5 days at 37 °C with gentle agitation. Organoids were subsequently washed extensively in PBST for 24 hours, changing the wash buffer every 4-6 hours. Secondary antibody incubation was performed under the same conditions as the primary antibody incubation, followed by another 24-hour wash in PBST. For refractive index matching, we utilized the CUBIC method. Immunolabeled organoids were immersed in 50% CUBIC-1 solution (diluted in water) overnight at 37 °C, followed by incubation in 100% CUBIC-1 solution for 2-3 days at 37 °C until transparency was achieved. The clearing solution was refreshed daily. Cleared organoids were mounted in chambers filled with the appropriate refractive index matching solution. Images were acquired using a LEICA STELLARIS 5 microscope to capture high-resolution, three-dimensional stacks. Three-dimensional reconstructions were performed using image analysis software with Fiji (ImageJ).

### RNA isolation and quantitative RT-PCR

Total RNA was extracted from cells or organoids using the Quick-RNA MicroPrep kit (ZymoResearch). RNA was subjected to quantitative real-time PCR in accordance with the protocol provided by one-step SYBR green RT-PCR Kit (Cwbio). The transcripts were quantitated and normalized to the internal GAPDH control. The primers used in the experiments are listed in **Table S2**. The PCR conditions were 1 cycle at 95 °C for 5 min, followed by 40 cycles at 95 °C for 15 s, 60 °C for 1 min, and 1 cycle at 95 °C for 15 s, 60 °C for 15 s, 95 °C for 15 s. The results were calculated using the 2^-△△CT^ method according to the GoTaq qPCR Master Mix (Promega) manufacturer’s specifications.

### Western blot analysis

H9^GTM^ cells treated by NAEK for 2 days were lysed in RIPA lysis buffer (Applygen) supplemented with complete protease inhibitor cocktail (Roche), and cell debris was removed by centrifugation at 4 °C. Protein was extracted with lysis buffer and quantified by BCA assay (Thermo). A total of 100 µg of protein from each sample was boiled with loading buffer, separated on 4-12% NuPAGE (Invitrogen), and electroblotted onto a polyvinylidene difluoride membrane. The membrane was blocked with 5% (v/v) non-fat milk in TBST (50 mM Tris-HCl, 150 mM NaCl and 0.02% Tween-20, pH 7.5) at room temperature for 1h and then incubated with rabbit or mouse polyclonal antibodies overnight at 4 °C. Anti-ETV2 antibodies (1:5,000, ab181847, Abcam) and GAPDH (1:50,000, 60004-1-Ig, Proteintech), were diluted in TBST containing 5% (v/v) of defatted milk. The membranes were rinsed three times with TBST and then incubated with horseradish peroxidase-conjugated goat anti-rabbit/mouse IgG (1:50,000) at room temperature for 1h. After extensive washing, the protein bands were developed using an enhanced chemiluminescent detection kit (Millipore). The bands were visualized in a ChampChemi®910 Chemiluminescent Imaging System (SINSAGE). Western blot experiments were performed at least twice in parallel and representative results were reported (**Supplementary Figure 22**).

### ZIKA virus infection

ZIKA virus (ZIKV) was gifted and tittered (1.0×10^7^ pfu/mL) by Anhui Medical University. vhCOs on day 90 were directly infected with ZIKV or Mock at the MOI of 0.5 for 24 h, and then replaced with fresh medium for continued culture for 6 days. After 7 days, vhCOs were collected for immunostaining and snRNA-seq. For immunostaining, vhCOs were stained with primary antibody listed in **Table S1**. For sc-RNA sequencing, vhCOs were quick frozen in liquid nitrogen for 1 min and then stored at -80 °C until performing snRNA-seq (Singleron Bio.). Overall size of organoids was measured under calibrated 4× bright field microscope.

### Flow cytometry

For flow cytometry analysis, the organoids were dissociated into single cells with Accutase, and then fixed with 4% PFA at room temperature for 20 minutes. Single cells were stained with Fluorescence labeled antibodies (**Table S1**) at RT for 30 minutes. After 3 times washing with 1% BSA in PBS, cells were analyzed using a BD FACSCelesta^TM^ flow cytometer. Single-color and isotype controls served as compensation and negative gating (**Supplementary Figure 23**).

### Whole-mount Immunostaining of organoids

We performed whole-mount immunostaining followed by confocal microscopy to examine the localization and organization of EC networks within the vhCOs. Organoids were washed with PBS and fixed overnight in 4% paraformaldehyde (PFA) at 4 °C. After washing the organoids with PBS for 3 times, the organoids were blocked overnight at RT in 0.5% BSA and 0.125% Triton-100 in PBS. Organoids were incubated in primary antibodies (anti-CD31, 1:200; anti-SOX2, 1:200) and diluted in 0.5% BSA and 0.125% Triton-100 in PBS for 2 d at 4 °C. Then, organoids were incubated with Alexa Fluor Dyes (1:500) for 2 h following nuclei staining with DAPI (1:1,000) for 1 h. After washing with PBS for 3 times, stained organoids were photographed under a LEICA STELLARIS 5 microscope.

### ELISA assay

vhCOs were treated with or without LPS (100 ng/mL) for 24 h, then collected the supernatant for inflammatory factors measurement. Anti-human IL6 and TNF-α ELISA kits (Elabscience, E-EL-H6156, E-EL-H0109, respectively) were used according to manufacturer’s protocol. Samples were analyzed in a multifunctional enzyme-linked instrument (Tecan INFINITE 200 PRO, Austria) with an optical density (OD) of 450 nm. Three batches of vhCOs were used for this experiment.

### TEER analysis

TEER in three-dimensional organoids was measured by placing micro-electrode probes in three different regions of the spheres. TEER measurements were performed by using an electrochemical impedance analyzer (Z100, eDAQ). For each control hCO and vhCO (days 30, 60 and 90, n = 3), three different measurements were performed and Z navigator was used to produce electrochemical impedance spectroscopy data from the organoids. ZMAN software (v.2.32), both metal and battery supercapacitor circuits, was used to analyze the data. Finally, resistivity is calculated as follows:

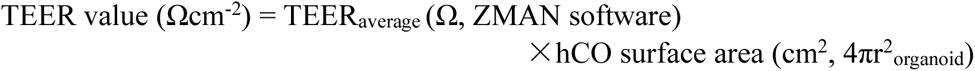

Where r represents the radius of the organoid.

### Single-cell nuclei isolation sorting

Organoids were harvested and were washed in pre-cooled PBSE (PBS buffer containing 2 mM EGTA). Nuclei isolation was carried out using GEXSCOPE® Nucleus Separation Solution (Singleron Biotechnologies, Nanjing, China) according to the manufacturer’s product manual. Isolated nuclei were resuspended in PBSE to 10^6^ nuclei per 400 µL, fltered through a 40 µm cell strainer, and counted with Trypan blue. Nuclei enriched in PBSE were stained with DAPI (1:1,000) (TermoFisher Scientifc, D1306). Nuiclei were defined as DAPI-positive singlets.

### Single nucleus RNA-sequencing library preparation

The concentration of single nucleus suspension was adjusted to 3-4 × 10^5^ nuclei/mL in PBS. Single nucleus suspension was then loaded onto a microfluidic chip (GEXSCOPE® Single NucleusRNA-seq Kit, Singleron Biotechnologies) and snRNA-seq libraries were constructed according to the manufacturer’s instructions (Singleron Biotechnologies). The resulting snRNA-seq libraries were sequenced on an Illumina novaseq 6000 instrument with 150 bp paired end reads.

### The analysis of snRNA-seq data

Cellranger software was used for mapping raw data to the human genome (version hg38 [v1.2.0]). Then, data were processed with Seurat (v5.1.0) under R (v4.3.3) environment. For quality control, cells expressing less than 300 genes or less than 1000 UMIs or more than 20% mitochondrial gene content were removed, and genes expressed in less than 10 cells were excluded for the following analysis. Putative doublets were removed using DoubletFinder (v2.0.4). Variance-stabilizing transformation (VST) was used for searching for the highly variable genes, and the top 2000 genes were chosen to do the downstream analysis. The cells were clustered and reduced into the UMAP space by PCA, with the principal components (PCs) selected based on ElbowPlot function. DEGs in each cluster were identified by more than 1.4-fold change and p-value<.05 with Wilcoxon rank-sum test. GO enrichment analysis was carried out by ‘ClusterProfiler’ and org.Hs.eg.db in R software program with false discovery rate (FDR) < 0.05 considered as statistical significance. Monocle (v2.32.0) package was used to analyze the developmental trajectory of cell types (using the recommended parameters). The dynamic model of scVelo (v0.3.3 based on Python 3.10) was used for modeling. Except for the vhCO+Z.V. group, the top 20% of genes with differential kinetic were used for correction. The cell differentiation trajectories were then visualized on the UMAP embedding. Cell-cell communication analysis was performed using Cellchat (v1.6.1) with default settings. Regulon expression was analyzed using pySCENIC (v0.12.1) with default settings. Based on the computed threshold, the open/off status of each regulon in each cell was binarized. The open proportion of each regulon was then calculated for each cell type. For GSVA, we transformed each cell cluster into a pseudobulk, used the post-log data and based on a Gaussian distribution to calculate enrichment scores. For the correlation between cell clusters, we selected the top 1000 genes with the greatest variability and calculated the Spearman correlation between clusters. To define the characteristic gene sets of brain regions while addressing the sparsity of scRNA-seq data, we first transformed Braun et al.’s scRNA-seq data into pseudobulk data according to brain regions and donors. Differential gene analysis was then performed using DESeq2. We employed two methods for identifying region-specific genes: 1, Binarized: Data from one brain region were fixed, while data from all other regions were averaged. A single comparison was sufficient to identify specific genes for a given region; 2, Paired: Differentially expressed genes (DEGs) were cyclically compared between a target brain region and all others, with the final gene set derived as the intersection of these comparisons. When setting a log2 fold-change (log2FC) threshold of 2, the binarized method yielded smaller but more specific gene sets. However, this strict threshold often resulted in empty sets for the paired method. To balance specificity and inclusiveness, we selected DEGs based on log2FC > 1 and FDR < 0.05. Additionally, we extracted a neuron-only sub-dataset and repeated the analysis. During GSVA enrichment, we excluded early brain samples (< PCW6) due to the limited number of specific genes detected in these stages.

### The reference mapping analysis based on deep generative model and transfer learning

Reference atlases from the following sources were used: Human fetal whole brain^[29]^, Human neocortex^[30]^, Human gastrula^[31]^, and Human vascular brain tissue^[40]^. Versions that had already been clustered and annotated by the original researchers were prioritized. Otherwise, manual annotation was performed. For mapping analysis, scVI and scANVI were used to train the reference atlas, prioritizing batch information when available, except for batches strongly correlated with age. Then, HNOCA-tools was used to retrain our atlas with the reference atlas. A weighted k-nearest neighbors (KNN) network (k = 100) was constructed to assign labels to each query cell, with the best label selected as the transferred annotation. Finally, the presence fraction of query dataset was visualized in the UMAP embedding of the reference atlas. Mapping analysis was performed for the entire altas as well as for subsets of each cell group.

### Subcutaneous implantation of vhCOs

hCOs and vhCOs grown for 90 d were embedded with Matrigel and used for implantation, and Matrigel served as negative control. Mice were placed into a chamber by providing 2% isoflurane for anesthetization. Then, mice were fixed in a laminar hood to cut a small incision at each back leg. Matrigel embedded hCOs and vhCOs were subcutaneously placed into the incision of the right and left back legs, respectively. Once organoids were inserted, the wound was closed with sutures and buprenorphine (0.1 mg/kg) was administered for pain relief. Subsequently, implanted organoids were analyzed via MRI analysis and immunofluorescence staining.

### FITC-dextran perfusion assay

After 30 d of organoid implantation, mice were injected with 15 mg/mL FITC dextran via a perfusion device. Briefly, the perfusion device was inserted in the left ventricle, and then we punctured the right ventricle. FITC-dextran was immediately injected into mice with a 5 mL/min flow rate via a peristaltic pump. After monitoring the color change in the blood vessels of mice (around 3-5 min), the mice brains, hCOs and vhCO were explanted and imaged for FITC via confocal microscopy. Then, explanted tissues were fixed and sliced for further immunofluorescence staining.

### Statistical analysis

All data were analyzed using the GraphPad Prism 9 software. For the statistical analysis of other results, statistical evaluation was performed by Student’s unpaired t-test or one-way ANOVA with Tukey’s multiple comparisons test. Data are presented as means ± SD or as described in the corresponding legends. A probability of p < 0.05 was considered as statistically significant. For annotations of significance, *p < 0.05; **p < 0.01; ***p < 0.001; ****p < 0.0001.

### Data availability

Single nucleus transcriptome data are available in the Gene Expression Omnibus under accession code GSE300436. All other data supporting this study are available within this paper and its Supplementary Information.

## Acknowledgements

This work was supported by National Key R&D Program of China (No. 2022YFA1104700), National Key Research and Development Program of China (No. 2024YFA1108000), the Space Application System of China Manned Space Program (KJZ-YY-NSM0505), and Noncommunicable Chronic Diseases-National Science and Technology Major Project (No. 2024ZD0531000).

The authors would like to acknowledge Y.Q. W., P.W. D., P. W., X. C. and W.P. Z., in the Biomedical Platform at Suzhou Institute for Advanced Research, University of Science and Technology of China, for technical support in device manufacturing. Some of the plasmids used in this work were kindly provided by L.Z D. from Peking University.

## Author contributions

J.H. Q. and H.S. L. conceived the idea and designed the experiments. H.S. L. performed the experiments and offered the original images. H.S. L. and Y.Q. W. organized the results, analyzed the data, prepared the figures and wrote the manuscript. H. D. is responsible for bioinformatics analysis. Y.Q. W., H. D., Y.X. Q., and P. W., edited the manuscript and gave technical advice. Zika virus was provided by Professor L. W. from Anhui Medical University. H.Y. Z. contributed to the revision of the manuscript. J.H. Q. supervised the study.

## References

1. M. A. Lancaster, M. Renner, C. A. Martin, et al., “Cerebral Organoids Model Human Brain Development and Microcephaly,” Nature 501, no. 7467 (2013): 373–379, 10.1038/nature12517.

2. X. Meng, D. Yao, K. Imaizumi, et al., “Assembloid CRISPR Screens Reveal Impact of Disease Genes in Human Neurodevelopment,” Nature 622, no. 7982 (2023): 359–366, 10.1038/s41586-023-06564-w.

3. Z. He, L. Dony, J. S. Fleck, et al., “An Integrated Transcriptomic Cell Atlas of Human Neural Organoids,” Nature 635 (2024): 690–698.

4. M. Wang, L. Zhang, S. W. Novak, et al., “Morphological Diversification and Functional Maturation of Human Astrocytes in Glia-Enriched Cortical Organoid Transplanted in Mouse Brain,” Morphological diversification and functional maturation of human astrocytes in glia-enriched cortical organoid transplanted in mouse brain, Vol. 43, 2024, 10.1038/s41587-024-02157-8.

5. K. Akassoglou, D. Davalos, A. S. Mendiola, et al., “Pioneering Discovery and Therapeutics at the Brain-Vascular-Immune Interface,” Cell 187, no. 21 (2024): 5871–5876.

6. C. N. Hall, C. Reynell, B. Gesslein, et al., “Capillary Pericytes Regulate Cerebral Blood Flow in Health and Disease,” Nature 508, no. 7494 (2014): 55–60.

7. R. D. Bell, E. A. Winkler, A. P. Sagare, et al., “Pericytes Control Key Neurovascular Functions and Neuronal Phenotype in the Adult Brain and during Brain Aging,” Neuron 68, no. 3 (2010): 409–427.

8. B. Cakir, Y. Tanaka, F. R. Kiral, et al., “Expression of the Transcription Factor PU.1 Induces the Generation of Microglia-like Cells in Human Cortical Organoids,” Nature Communications 13, no. 1 (2022): 1–15, 10.1038/s41467-022-28043-y.

9. J.-A. L. Kelsey C. Martin Mhatre V. Ho, Ruben Martin and Stephen L. Buchwald, J.-A. L. Mhatre V. Ho and Kelsey C. Martin, C. Craik, A. Manuscript, and Kantrowitz, “Microglia Promote Learning-Dependent Synapse Formation through BDNF,” Cell 155, no. 7 (2013): 1596– 1609, 10.1016/j.cell.2013.11.030.Microglia.

10. Y. Shi, L. Sun, M. Wang, et al., “Vascularized Human Cortical Organoids (VOrganoids) Model Cortical Development in Vivo,” PLoS Biology 18, no. 5 (2020): 1–29, 10.1371/journal.pbio.3000705.

11. T. Wang, B. D. Gastfriend, V. Mcdonald, J. P. Steiner, A. G. Elkahloun, and A. Nath, “Derivation of a Human Brain Organoid with Microglia Development,” no. January (2025), 10.3791/67491.

12. X. Y. Sun, X. C. Ju, Y. Li, et al., “Generation of Vascularized Brain Organoids to Study Neurovascular Interactions,” eLife 11 (2022): 1–28, 10.7554/eLife.76707.

13. B. Cakir, Y. Xiang, Y. Tanaka, et al., “Engineering of Human Brain Organoids with a Functional Vascular-like System,” Nature Methods 16, no. 11 (2019): 1169–1175, 10.1038/s41592-019-0586-5.

14. J. W. Chin, “Expanding and Reprogramming the Genetic Code of Cells and Animals,” Annual Review of Biochemistry 83 (2014): 379–408, 10.1146/annurev-biochem-060713-035737.

15. D. D. Young, and P. G. Schultz, “Playing with the Molecules of Life,” ACS Chemical Biology 13, no. 4 (2018): 854–870, 10.1021/acschembio.7b00974.

16. J. C. W. Willis, and J. W. Chin, “Mutually Orthogonal Pyrrolysyl-TRNA Synthetase/TRNA Pairs,” Nature Chemistry 10, no. 8 (2018): 831–837.

17. S. Lee, C. Park, J. W. Han, et al., “Direct Reprogramming of Human Dermal Fibroblasts into Endothelial Cells Using ER71/ETV2,” Circulation Research 120, no. 5 (2017): 848–861, 10.1161/CIRCRESAHA.116.309833.

18. K. Wang, R. Z. Lin, X. Hong, et al., “Robust Differentiation of Human Pluripotent Stem Cells into Endothelial Cells via Temporal Modulation of ETV2 with Modified MRNA,” Science Advances 6, no. 30 (2020): 1–15, 10.1126/sciadv.aba7606.

19. A. H. M. Ng, P. Khoshakhlagh, J. E. Rojo Arias, et al., “A Comprehensive Library of Human Transcription Factors for Cell Fate Engineering,” Nature Biotechnology 39, no. 4 (2021): 510– 519, 10.1038/s41587-020-0742-6.

20. M. A. Lancaster, and J. A. Knoblich, “Generation of Cerebral Organoids from Human Pluripotent Stem Cells,” Nature Protocols 9, no. 10 (2014): 2329–2340, 10.1038/nprot.2014.158.

21. D. H. Lee, T. M. Kim, J. K. Kim, and C. Park, “ETV2/ER71 Transcription Factor as a Therapeutic Vehicle for Cardiovascular Disease,” Theranostics 9, no. 19 (2019): 5694–5705, 10.7150/thno.35300.

22. T. M. Kim, R. H. Lee, M. S. Kim, C. A. Lewis, and C. Park, “ETV2/ER71, the Key Factor Leading the Paths to Vascular Regeneration and Angiogenic Reprogramming,” Stem Cell Research and Therapy 14, no. 1 (2023): 1–16, 10.1186/s13287-023-03267-x.

23. W. Gong, S. Das, J. E. Sierra-Pagan, et al., “ETV2 Functions as a Pioneer Factor to Regulate and Reprogram the Endothelial Lineage,” Nature Cell Biology 24, no. 5 (2022): 672–684, 10.1038/s41556-022-00901-3.

24. S. Y. Oh, J. Y. Kim, and C. Park, “The ETS Factor, ETV2: A Master Regulator for Vascular Endothelial Cell Development,” Molecules and Cells 38, no. 12 (2015): 1029–1036, 10.14348/molcells.2015.0331.

25. N. Koyano-Nakagawa, and D. J. Garry, “Etv2 as an Essential Regulator of Mesodermal Lineage Development,” Cardiovascular Research 113, no. 11 (2017): 1294–1306, 10.1093/cvr/cvx133.

26. M. S. Kim, R. Lee, D. H. Lee, et al., “ETV2/ER71 Regulates Hematovascular Lineage Generation and Vascularization through an H3K9 Demethylase, KDM4A,” iScience 28, no. 1 (2025): 111538, 10.1016/j.isci.2024.111538.

27. T. Lazarov, S. Juarez-Carreño, N. Cox, and F. Geissmann, “Physiology and Diseases of Tissue-Resident Macrophages,” Nature 618, no. 7966 (2023): 698–707.

28. E. G. Perdiguero, K. Klapproth, C. Schulz, et al., “Tissue-Resident Macrophages Originate from Yolk-Sac-Derived Erythro-Myeloid Progenitors,” Nature 518, no. 7540 (2015): 547–551.

29. E. Braun, M. Danan-Gotthold, L. E. Borm, et al., “Comprehensive Cell Atlas of the First-Trimester Developing Human Brain,” Science 382, no. 6667 (2023), 10.1126/science.adf1226.

30. L. Wang, C. Wang, J. A. Moriano, et al., “Molecular and Cellular Dynamics of the Developing Human Neocortex at Single-Cell Resolution,” Molecular and cellular dynamics of the developing human neocortex at single-cell resolution, 2024, 10.1038/s41586-024-08351-7.

31. B. Zeng, Z. Liu, Y. Lu, et al., “The Single-Cell and Spatial Transcriptional Landscape of Human Gastrulation and Early Brain Development,” Cell Stem Cell 30, no. 6 (2023): 851–866.e7, 10.1016/j.stem.2023.04.016.

32. X. Qian, H. N. Nguyen, M. M. Song, et al., “Brain-Region-Specific Organoids Using Mini-Bioreactors for Modeling ZIKV Exposure,” Cell 165, no. 5 (2016): 1238–1254, 10.1016/j.cell.2016.04.032.

33. P. P. Garcez, E. C. Loiola, R. M. Da Costa, et al., “Zika Virus: Zika Virus Impairs Growth in Human Neurospheres and Brain Organoids,” Science 352, no. 6287 (2016): 816–818, 10.1126/science.aaf6116.

34. J. Dang, S. K. Tiwari, G. Lichinchi, et al., “Zika Virus Depletes Neural Progenitors in Human Cerebral Organoids through Activation of the Innate Immune Receptor TLR3,” Cell Stem Cell 19, no. 2 (2016): 258–265, 10.1016/j.stem.2016.04.014.

35. M. Watanabe, J. E. Buth, N. Vishlaghi, et al., “Self-Organized Cerebral Organoids with Human Specific Features Predict Effective Drugs to Combat Zika Virus Infection,” 21, no. 2 (2017): 517–532, 10.1016/j.celrep.2017.09.047.Self-organized.

36. E. M. Abud, R. N. Ramirez, E. S. Martinez, et al., “NeuroResource,” Neuron 94, no. 2 (2017): 278–293.e9, 10.1016/j.neuron.2017.03.042.

37. P. R. Ormel, R. V. De Sá, E. J. Van Bodegraven, et al., “Microglia Innately Develop within Cerebral Organoids,” (2018), 10.1038/s41467-018-06684-2.

38. G. Popova, S. S. Soliman, C. N. Kim, et al., “Human Microglia States Are Conserved across Experimental Models and Regulate Neural Stem Cell Responses in Chimeric Organoids,” Cell Stem Cell 28, no. 12 (2021): 2153–2166.

39. D. P. Nguyen, H. Lusic, H. Neumann, P. B. Kapadnis, A. Deiters, and J. W. Chin, “Genetic Encoding and Labeling of Aliphatic Azides and Alkynes in Recombinant Proteins via a Pyrrolysyl-TRNA Synthetase/TRNACUA Pair and Click Chemistry,” Journal of the American Chemical Society 131, no. 25 (2009): 8720–8721, 10.1021/ja900553w.

40. T. Wälchli, J. Bisschop, P. Carmeliet, et al., “Shaping the Brain Vasculature in Development and Disease in the Single-Cell Era,” Nature Reviews Neuroscience 24, no. 5 (2023): 271–298.

